# Stability and affinity evaluation of aptamers as a biorecognition system for testosterone and its analogs

**DOI:** 10.1101/2024.03.18.585618

**Authors:** Ariadna Medina-Benítez, Claudia A. García-Alonso, Roberto Fragoso-Rodríguez, Ana Laura Torres-Huerta, Aurora Antonio-Pérez

**Affiliations:** Escuela de Ingeniería y Ciencias, Tecnológico de Monterrey, Campus Estado de México, Av. Lago de Guadalupe KM 3.5, Margarita Maza de Juárez, Ciudad López Mateos, Atizapán de Zaragoza 52926, Estado de México, Mexico. / /

**Keywords:** biosensor, aptasensor, *in silico* analysis, molecular docking, aptamer, testosterone

## Abstract

The present project results from the need to develop a biosensor for the detection of testosterone and synthetic analogs in nutritional supplements. A biosensor is composed of two systems: (1) the biorecognition system and (2) the transducer system. We propose the use of aptamers as a biorecognition system, given their functional capacities to recognize and “capture” specific small analytes. For this study, eight aptamers sequences with previous reports of interaction with testosterone were evaluated to select the best candidate on regard with the best affinity and structural stability at different ionic concentrations and temperature values. *In silico* analysis involves predicting secondary and tertiary structures in different conditions and molecular docking. *In vitro* analysis. tracks the changes in the folded and unfolded aptamer dimensions monitored by dynamic light scattering under the previously *in silico* conditions, and performs relative capture tests with testosterone and testosterone undecanoate by immobilized aptamers on gold nanoparticles. The outcomes of the *in silico* approach aligned with the experimental data collected, exhibit that the nucleotide composition and aptamers binding structure, intervene in their affinity and biosensor functionality. The aptamers apT5 and P4G13 were discarded as possible candidates because they did not show stability in experimental conditions (temperature and ion plug). TESS2, TESS3, T4, T5.1, and T6 proved to be functional aptamers because they showed good affinity with the three analytes, however, TESS1 turned out to be the best candidate because of its stability and number of interactions, which translates into a greater affinity, compared to the rest.

## 1. Introduction

The use of nutritional supplements has spread massively over time. According to Daher, et al (2022), the prevalence of consumption of nutritional supplements among athletes in different studies varied from 11 to 100% of the participants, being performance improvement the main reason [1]. Nonetheless, there are currently no strict regulations that govern the purity and effectiveness of nutritional supplements. Some reports indicate that dietary supplements can be exposed to the presence, high concentration, and non-specific composition of androgens, which represent a risk factor for the development of certain diseases, endangering the health of those who consume them [2,3]. Testosterone and/or its synthetic variables can interact with the components of the supplement androstenedione and lead to metabolic alterations, serious health effects, and addiction due to long-term consumption [4 - 6]. This scenario points out the need to develop effective tools for detecting these compounds in dietary supplements, such as biosensors.

Biosensors are devices that allow the recognition of an analyte of interest using a signal transduction system and a biological system and are used in several areas of application due to their versatility, easy application, and ability to provide quantitative and qualitative information. An aptasensor is a biosensor that employs aptamers for the biorecognition of the analyte of interest. Aptamers are defined as a simple sequence of oligonucleotides (generally fewer than 100) with an average molecular weight of 5 - 25 kDa, that adopt different tertiary structure configurations to selectively bind their ligand (the analyte), through electrostatic interactions, hydrogen bonds, bonds between aromatic rings, interactions between the nucleobases of the aptamer, and complementarity with the three-dimensional structure [7,8].

Their high specificity is owed to the high flexibility and torsion angles between the (deoxy)ribose-phosphodiester bonds of the chain. The most common technique for aptamer development is the Systematic Evolution of Ligands by Exponential Enrichment (SELEX), a repetitive procedure in which the target is consistently exposed to a collection of random oligonucleotides, leading to the progressive enrichment of sequences that exhibit the strongest binding affinity towards the analyte. Consequently, the selected aptamers possess high selectivity and affinity, characterized by dissociation constants (Kd) ranging from picomolar to nanomolar intervals (pM–μM), regardless of the nature and size of the target [8].

However, these features are subjected to structural changes derived from temperature and ionic concentration [8,9]. Ionic concentration has been shown to have a direct impact on aptamer conformation, as well as on their binding affinity, decreasing both by either the shielding of DNA or of the ligand’s negative charges by sodium ions [11,13], whereas temperature has a higher impact to the binding affinity and constantly their evaluation its related to the future application [12]. Therefore, evaluating the secondary and tertiary structures and the changes in them to foresee the conditions that lead to the best aptamer-ligand interaction and optimization of the aptasensor design is crucial [9,10,14,15]. *In silico* analysis is based on the modeling of secondary and tertiary structures from a simple chain sequence (primary structure), by the application of computational methods based on mathematical models, to understand the mechanisms of aptamer-ligand interactions. Nowadays there are websites and programs available for modeling structures that consider characteristics such as hydrophilicity, hydrophobicity, entropy (*S*), and enthalpy (*H*), and structural properties like inclination, bending, rise, slicing, shifting, and more [10,16], as well as for the molecular docking of proteins with their ligands, in order to analyze their interactions [7,10]. Thus, the objective of this project is to evaluate the interactions between different aptamers and androgens testosterone, androstenedione, and testosterone undecanoate, for the design of an aptasensor that enables their detection in dietary supplements.

In summary, the aptasensor is an alternative to the need to establish a method for detecting testosterone or its synthetic analogs in dietary supplements. The outlined features of the aptamers are translated into attributes of the aptasensor, fulfilling the prerequisites for the future development of a biosensor, but these features also respond to physicochemical conditions that must be analyzed.

*In silico* strategy in the present project follows three main steps: The first step is “Prediction of secondary structure”, in which we modify the parameters (temperature and ionic conditions) that command structure, folding, and stability, which remain highly dependent on the following steps. The second step is, “Prediction, and optimization of tertiary structure”, based on the secondary structure this step displays the structure and dynamics that include the intra-aptamer bindings as a unique 3D conformation and their respective arrangements like stems, hairpins, loops, pseudoknots, bulges, and/or G-quadruplexes. The third step, “Molecular docking” is to identify the potential binding sites and evidence of the aptamer-analyte complex bonding, also considering the structure and flexibility of the aptamer. The complete evaluation of each aptamer gives a more approach knowledge about their behavior, and response and permits it to be evaluated and compared with the other aptamers in the aim of selecting the best candidate [15,16].

Experimental studies were conducted to monitor the structural changes associated with the folding or unfolding driven by the exposure of the aptamers to the established temperature and chemical conditions of the *in silico* strategy. These samples were monitored by dynamic light scattering (DLS) to compare the predicted aptamer structural stability with experimental data. On the other hand, relative capture assays using testosterone and testosterone undecanoate were used to evaluate the molecular docking results. The aptamers in the nanoparticles must be immobilized for these capture tests to be conducted. In this document, it can be seen that through both experimental strategies applied in this study, it was possible to validate the predictive scope of the *in silico* analysis strategy developed [17].

## 2. Materials and methods

### 2.1. In silico analysis

The *in silico* methodology followed in this project is described according to the following flow chart, Figure 1.

**Figure 1.**
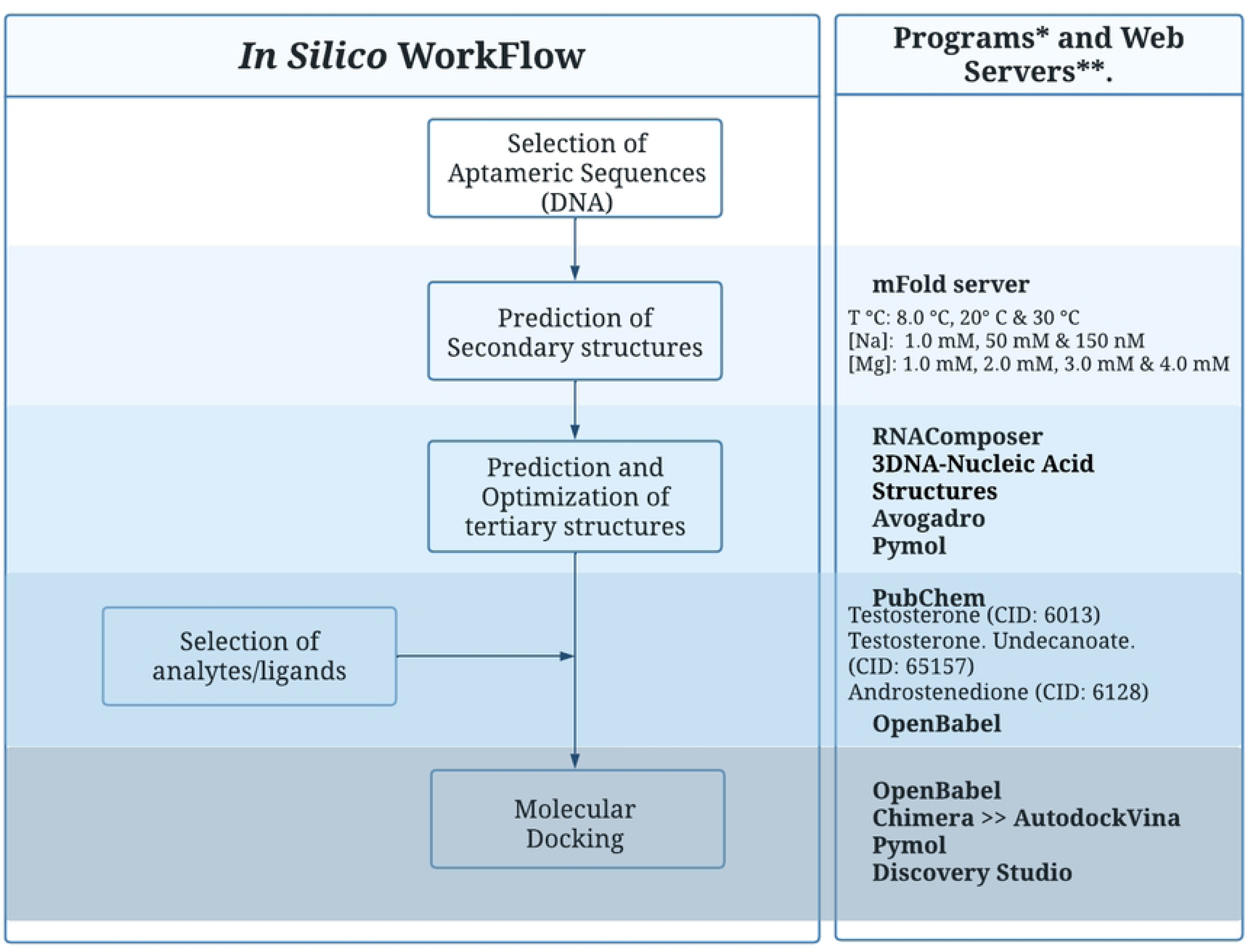
Flowchart, *in silico* methodology. * Programs: Avogadro (Avogadro: an open-source molecular builder and visualization tool. Version 1.0. Baltimore. http://avogadro.cc/) [32]. PyMOL (The PyMOL Molecular Graphics System, Version 2.0 Schrödinger, LLC) [25]. Open Babel (Open Babel: An open Chemical toolbox. Version 3.1.1, Arizona) [33]. Chimera (UCSF Chimera, production version 1.16 (build 42360)2021. Platform:Darwin64. Windowing system:aqua, California) [34]. Discovery Studio (BIOVIA, Dassault Systèmes, Discovery Studio Visualizer, v 21.1.0.20298, San Diego) [28]. ** Web - Servers: mFold server (http://www.unafold.org/mfold/applications/dna-folding-form.php). [22] RNAComposer (https://rnacomposer.cs.put.poznan.pl) [35,36]. PubChem (https://pubchem.ncbi.nlm.nih.gov) [26]. 3DNA-Nucleic Acid Structures web server (http://web.x3dna.org) [37].

#### 2.1.1. Selection of Aptameric Sequences

We select eight aptamer sequences that, according to the reported literature, are designed to detect testosterone and some of its synthetic analogs, also they present proven interaction activity; the name ID, sequence, and reference of each of the aptamers are specified, in Table 1.

**Table 1.**
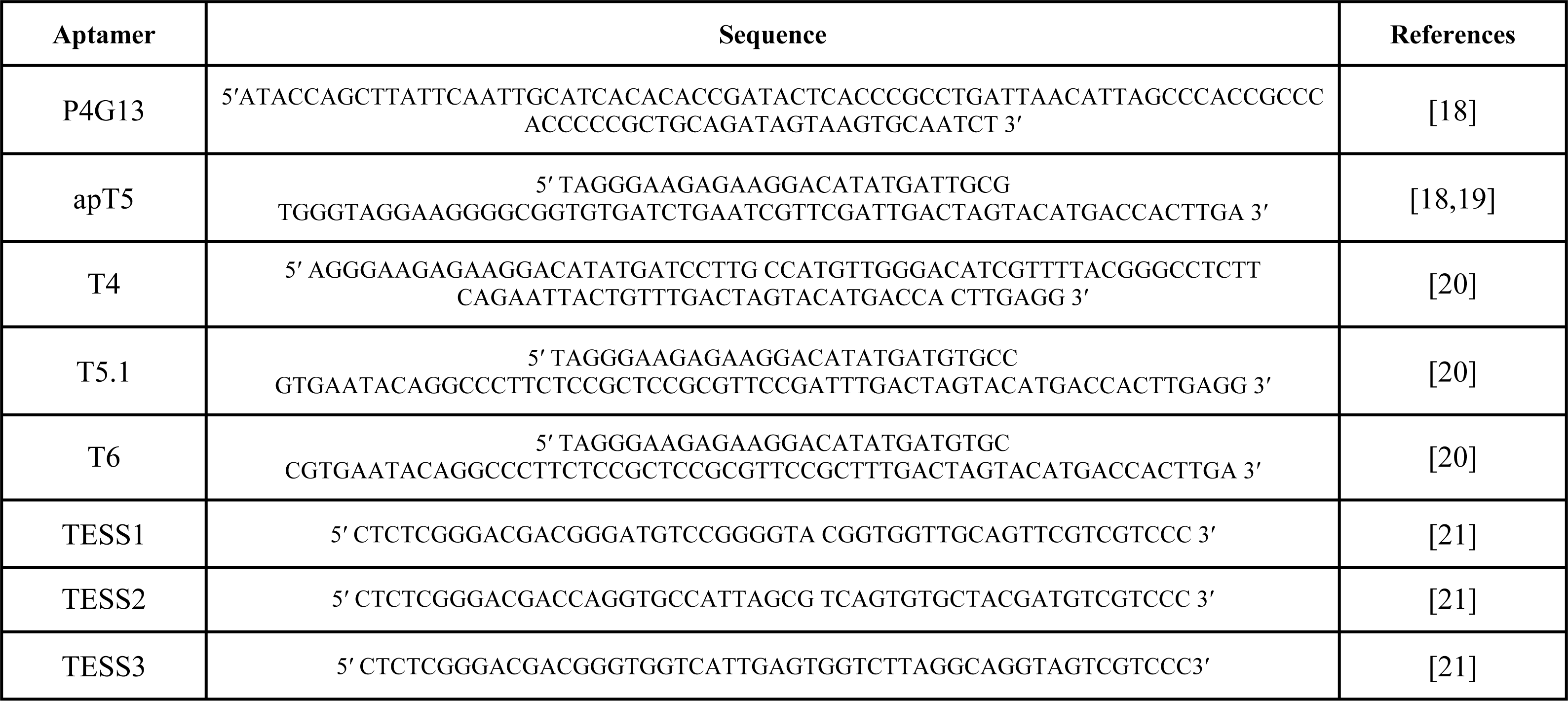
Aptamers are designed for the detection of testosterone and its synthetic analogs.

#### 2.1.2. Prediction of the Secondary Structure

For *in silico* analysis of the aptamer’s stability presented in Table A.1, the secondary structures of each sequence were modeled, individually modifying the following parameters:

● Folding temperature at 8°C, 20°C, and 30°C.
● Ionic plug:
○ Sodium concentration [Na^+^] at 1 mM, 50 mM, and 150 mM.
○ Magnesium concentration [Mg^2+^] at 1 mM, 2 mM, 3 mM, and 4 mM.

The library of these secondary structures was generated using the web platform mfold server, resulting in files of a CT format. Mfold server has been used extensively in publications concerning the generation of secondary structures. In general, the algorithm determines multiple optimal and suboptimal secondary structures based on the minimization of the free energy of Gibbs, ΔG, as well as the free energies contained in any base pair. Also folding and hybridization parameters have been measured *in vivo* or by laboratories before and after being incorporated into the mfold package [10,16,20,22].

#### 2.1.3. Prediction and Optimization of the Tertiary Structure

Tertiary structures were modeled for each child structure generated using the RNAComposer web server, where the CT format was converted to the dotted bracket format, and on the same server, the tertiary structure was obtained in “.pdb” format (Protein Databank File) [35,36]. Subsequently, the 3DNA-Nucleic Acid Structures web server was used to convert the structures from RNA to DNA [37]. The optimization of each of the structures was carried out with the Avogadro program to be visualized later with the PyMOL program [32,25]. The RNAComposer is also very often used software in publications that develop tertiary structures, since their *in silico* approach has proven to be identical to experimentation results, especially for hairpin and small structures. It is based on the automatic translation and a comparative relationship of the secondary structure fragments and the elements of a tertiary structure taking as reference the database "RNA FRABASE", where the structures with the higher negative value of free energy are used for the 3D structure [10,15,35,36].

#### 2.1.4. Selection of analytes/ligands

We obtained the tertiary structures of the analytes/ligands: testosterone (CID: 6013), testosterone undecanoate (CID: 65157), and androstenedione (CID: 6128) in PubChem web server in “.sdf” format (Simulation Description Format) and changed to “.pdb” using the Open Babel program [26,33].

#### 2.1.5. Molecular Docking

Molecular docking was carried out individually for each aptamer tertiary structure modeled and for each analyte/ligand using the AutoDock Vina tool in the Chimera program [27,34]. The generated files were modified with Open Babel software (“.pdbqt” to “.pdb” format) to visualize the interactions and links of union between the aptamer and target using PyMOL software and Discovery Studio [25,28,33]. AutoDock Vina algorithm is based on ΔG energy and empirical scoring that use a model that evaluates two kinds of data: the knowledge of the binding affinity for protein-ligand complexes, and the binding energy for each interaction element [10,16,27].

### 2.2. In vitro analysis

The *in vitro* methodology followed in this project is described according to the following flow chart, Figure 2.

**Figure 2.**
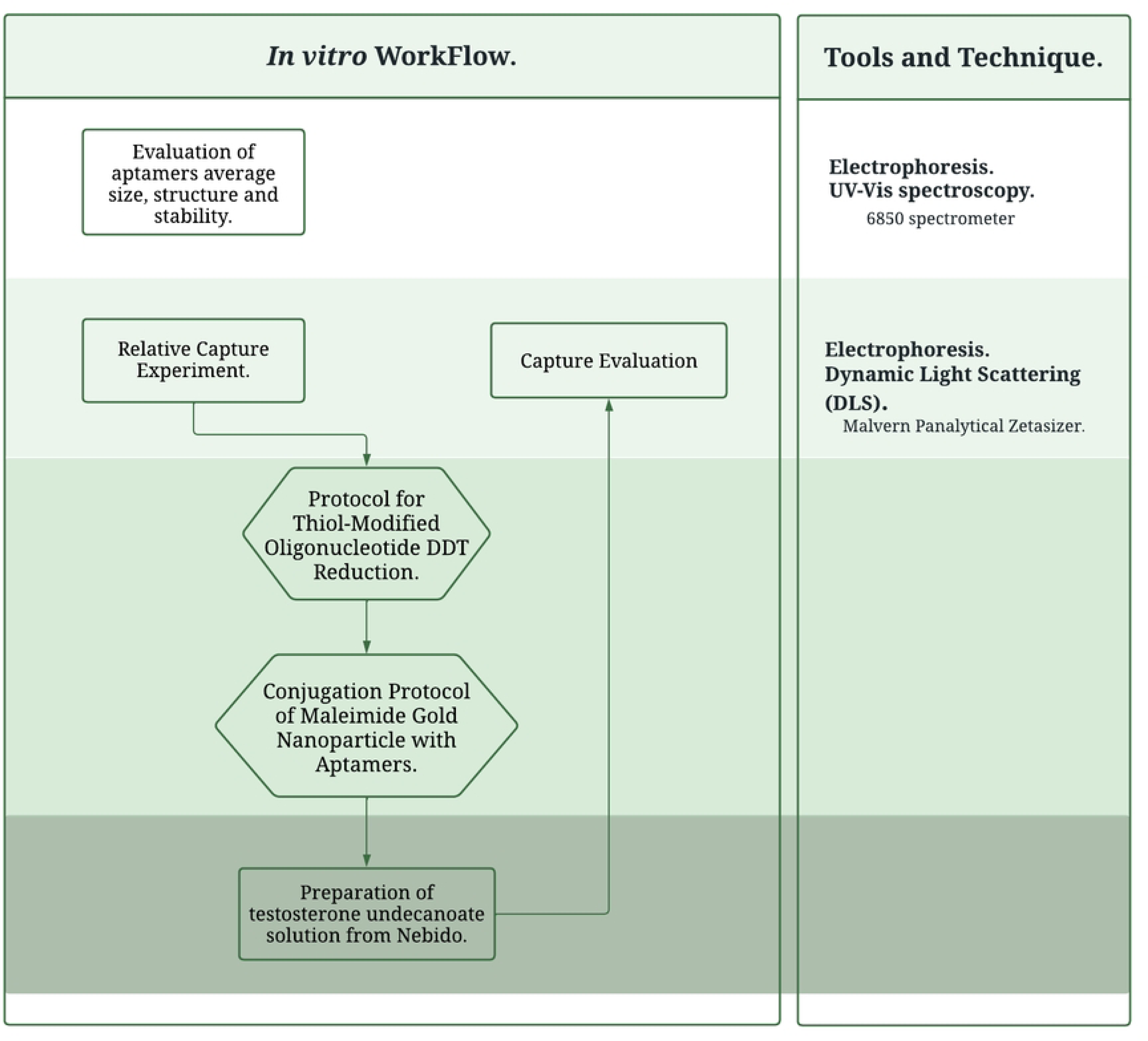
Flowchart, *in vitro* methodology.

#### 2.2.1. Evaluation of aptamers’ average size, structure, and stability

Aptamers T5, T6, TESS1, and TESS3 were resuspended with ddH2O with the corresponding volume to get a concentration of 100 μM, according to their manufacturing instructions. For the worked solutions: (1) we prepared 1:20 control dissolutions with dH2O to a total volume of 1 ml of each aptamer. (2) we prepared 1:50 dissolutions with dH2O to a total volume of 1 ml of each aptamer. (3) we prepared 1:50 dissolutions with a solution of NaCl at 1 mM, 50 mM, and 150 mM (individually) to a total volume of 1 ml of each aptamer. (4) we prepared 1:50 dissolutions with a solution of MgSO_4_ at 1 mM, 2 mM, 3 mM, and 4 mM (individually) to a total volume of 1 ml of each aptamer. Gel electrophoresis was performed as a general visualizer by protocol agarose gel at 3 %. Dynamic Light Scattering (DLS) technique was used to get the molecular weight (average size) of aptamers, 1 ml of worked solutions are displayed into cuvettes and analyzed with Malvern Panalytical Zetasizer using 2.0.0.98 software version and the parameters of 1.58 for material (DNA-aptamer) refractive index and zero for material absorption, and as the dispersant, we employed water with a refractive index of 1.33. Worked solutions (1) are directly evaluated at room temperature, (2) are displayed at the different folding temperatures (8°C, 20°C, and 30°C), while (3) and (4) are directly evaluated at room temperature.

#### 2.2.2. Relative capture experiment

As an immobilization strategy and possible adaptable system for the detection of testosterone and analogs, we follow the Protocol for Thiol-Modified Aptamers DDT Reduction from Sigma Aldrich and Cytiva and the Conjugation Protocol of Maleimide Gold Nanoparticles with Aptamers from Cytodiagnostics. The conjugation of aptamers T5.1, T6, TESS2, and TESS3 with AuNPs allows us to physically separate the aptamers with the analytical phase. The final conjugation solutions already have the sodium ion concentration (50 mM) and are displayed at room temperature (∼20 - 25 °C).

##### 2.2.2.1. Preparation of analytes

For testosterone: 80 μL of “Low Tiyel” were diluted in 4 mL of methanol and sonicated for 20 min, subsequently filtered through 0.45 μm nylon filters (Puradisc Sigma Aldrich). Then it was added 2 mL of saturated NaCl solution and centrifuged at 8 000 rpm for 20 min. Lastly, we filtered the supernatant with a 0.22 μm nylon filter (Puradisc Sigma Aldrich). The same step is followed to perform a calibration curve varying the amount (μL) of “Low Tiyel” and with the Low Tiyel polymer excipient as a blanch at Scan Spectrum Mode using the Jenway 6850 spectrometer.

For testosterone undecanoate: 40 μL of “Nebido” were diluted in 4 mL of methanol and sonicated for 20 min, subsequently filtered through 0.22 μm nylon filters (Puradisc Sigma Aldrich) by duplicate. The same step is followed to perform a calibration curve varying the amount (μL) of “Nebido” and with the Nebido oil excipient as a blanch at Scan Spectrum Mode using the Jenway 6850 spectrometer.

##### 2.2.2.2. Capture Evaluation

The formulations were prepared in adherence to the established protocol, comprising 5 μl of the biosensor system (30 μM) and 995 ml of the previously prepared testosterone solution, denoted as "Low Tiyel" (2.28 mM) and testosterone undecanoate solution, denoted as "Nebido" (4.77 mM). The biosensor system differs by the aptamer (1) T5.1, (2) T6, (3) TESS1, (4) TESS2 and (5) TESS3. All solutions were meticulously prepared with duplicate counterparts to ensure experimental reliability. Subsequently, the solutions underwent an incubation period of 30 minutes at room temperature on a shaker. The duration and analyte concentration (testosterone undecanoate solution) were selected at their maximum range to optimize binding events. Gel electrophoresis was performed as a general visualizer by the protocol of Agarose Gel at 1.2 %. Following the incubation period, centrifugation was executed at 13,000 rpm for 40 minutes across all biosensor systems, resulting in the capture of testosterone undecanoate in a pellet. The supernatant was carefully extracted for absorbance readings at 230 nm. This analysis was performed on a single quartz cell (1 mL) utilizing the Jenway 6850 spectrometer, ensuring precision in the measurement process.

## 3. Results

To demonstrate the effectiveness of our interaction modeling strategy, we conducted a literature review, searching for aptamers that were designed to bind different steroids. The selected aptamers, based on literature findings [18 – 21] were: apT5, P4G13, TESS1, TESS2, TESS3, T4, T5.1, and T6. They have based on reports of sequences with proven interaction capacity for testosterone.

The *in silico* molecular analysis started with DNA secondary structural prediction as a prerequisite to determine the tertiary structure and perform the molecular docking. Hybridization and melting temperatures were based on the free energy minimization algorithm and thermodynamics base pairing by Turner using the mfold server, since it’s suitable for RNA and DNA sequences has very close experimental prediction (83 %), and also mfold is more friendly to use, and to modify the conditions of temperature and ion plug than others [16,22,23]. We modified the parameters and obtained the aptamer model of each candidate varying the experimental conditions of ion plug and temperature, with the aim of evaluating their physicochemical stability. The modulation of the secondary structure displayed a ΔG value that represented the thermodynamic stability of the aptamer: the lower the value of ΔG, the more stable the structure is [10]. Table A.1 and Table A.2 contain the ΔG values that were obtained, and these values are visually represented in Figure 3. The evaluation of physicochemical stability for each aptamer is presented in Figure 3. The analysis was carried out at three different temperatures to evaluate simultaneously with ionic conditions and indicate with an arrow the aptamers that present a uniform behavior and greater stability. In Figure 3(a), the more stable aptamer was TESS1 (light blue color), having a bigger size and higher negative value of ΔG at any change of temperature and sodium concentration. By the modification of sodium ion (Na+), ΔG values ranged from 0.13 kcal/mol to -17.34 kcal/mol. For Figure 3(b), the ΔG value presents a clearer relation between the increase in magnesium concentration and the decrease in temperature, with ΔG values ranging from 0.59 kcal/mol to – 18.05 kcal/mol. TESS1 (light blue color) stands out as the most stable aptamer, followed by T6 (green color); also, TESS2 (red color) and TESS3 (dark blue color) presented a marked and consistent behavior.

**Figure 3.**
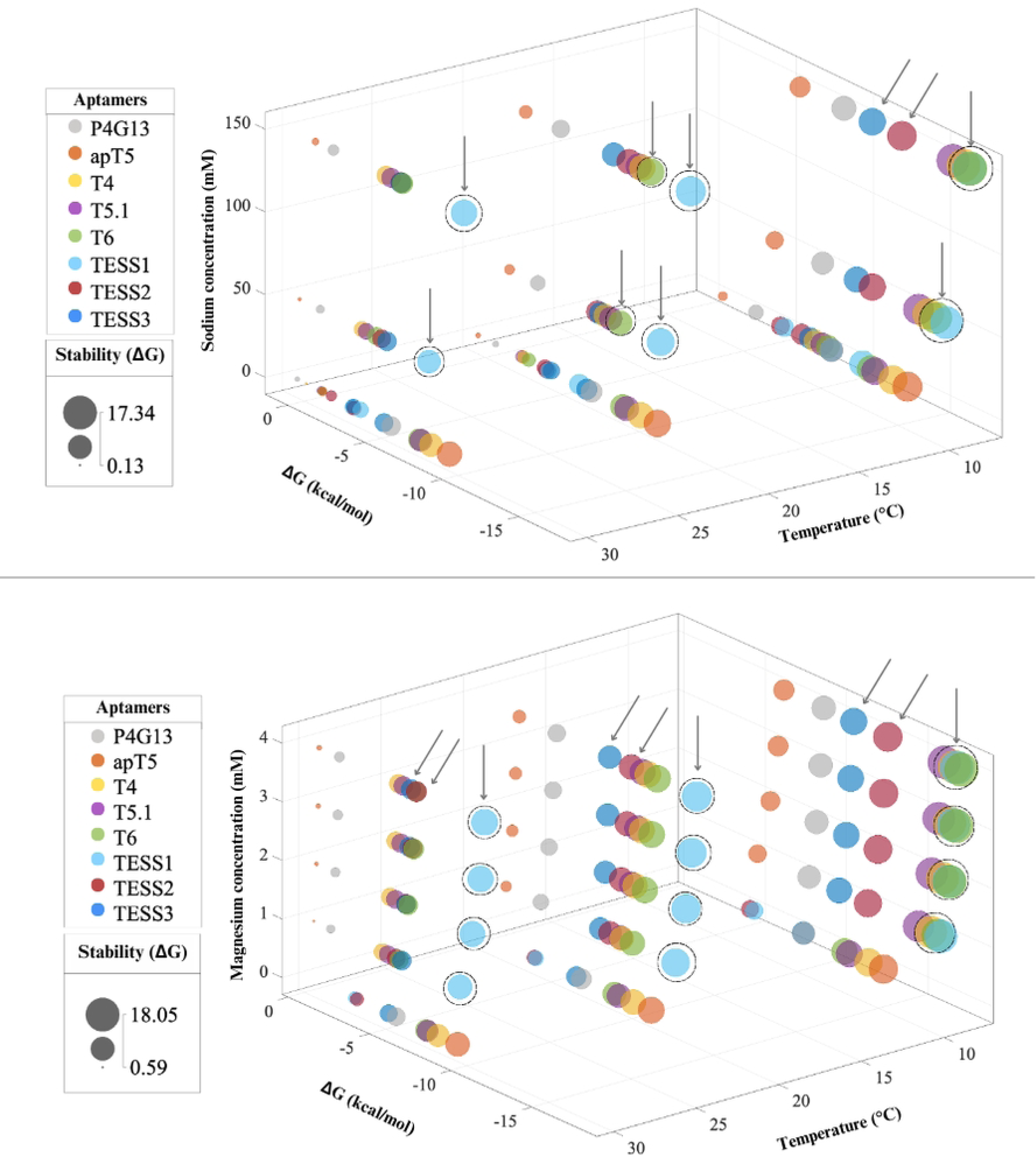
Bubble charts, for physicochemical stability evaluation. **(a)** Chart according to ΔG values from Table A.1, from the secondary structure prediction at different temperatures (8°C, 20°C, and 30°C) and sodium concentrations (0 mM, 1 mM, 50 mM, and 150 mM), where the size of each bubble is proportional to their negative ΔG values, in such a way that the bigger bubbles represent greater stability, under the criteria that locate them in the graph. **(b)** Chart according to ΔG values Table A.2, from the secondary structure prediction at different temperatures (8°C, 20°C, and 30°C) and magnesium concentration (0 mM, 1 mM, 2 mM, 3 mM, and 4 mM), where the size of each bubble is proportional to their negative ΔG values, in such a way that the larger bubbles represent greater stability, under the criteria that locate them in the graph. Charts developed on MATLAB: Optimization Toolbox version: 9.4 (R2022b), Natick, Massachusetts [24].

The significance of sodium and magnesium ions as adjustable parameters on the structural conformation of aptamers is based on the quantity of charged ions since their concentration impacts the formation of stable complexes. While the polarity of the charge determines the strength and stability of the bonds. Both ions are positively charged and are the most common present ions on worked solutions of aptamers, specifically, our future experimentations include the presence of sodium ion at a concentration between 50 mM to 150 mM, the main reason why the *in silico* analysis centers on cations evaluation. The difference between sodium ion and magnesium ion remains in the interaction intensity (monovalent and divalent respectively) between the negative charges of the aptamer that consequently interfere with their stability and 3D conformation.

As a continuing step for the *in silico* analysis, we modeled the tertiary structure from the secondary structure of the DNA sequences, intending to have a quantitative and a qualitative perspective for the behavior and stability of the eight proposed sequences in the current project, we present Figure 4.

**Figure 4.**
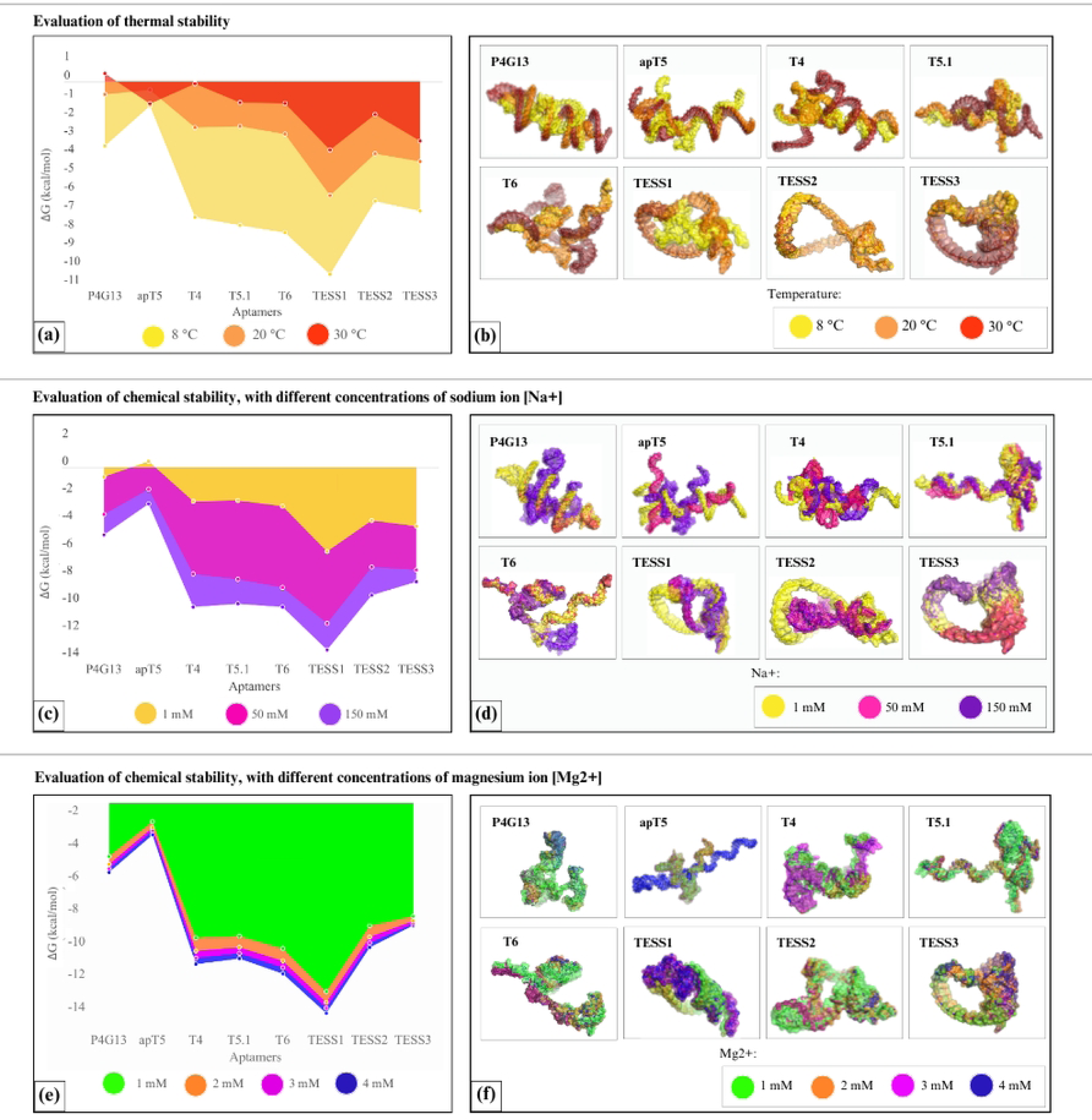
Evaluation of physicochemical stability. **(a)** Surface graph, according to ΔG values from **Table A.1**., from the secondary structure prediction at different temperatures with color correspondence to 8°C in yellow, 20°C in orange, and 30°C in red. Concentrations of sodium ion [Na+] and magnesium ion [Mg+2] were defined by the mFold server [22], up to the minimum permitted concentrations (0.01 M and 0.1 mM respectively). **(b)** Tertiary structures of the eight aptamers were modeled at different temperatures with the same color correspondence. **(c)** Surface graph, according to ΔG values from Table A.1., from the secondary structure prediction at different concentrations of sodium ion [Na+], with color correspondence to 1 mM in yellow, 50 mM in rosy color, and 150 mM in purple; with 20°C as constant temperature. **(d)** Tertiary structures of the eight aptamers modeled at different concentrations of sodium ion [Na+], with the same color correspondence. **(e)** Surface graph, according to ΔG values from Table A.1., from the secondary structure prediction at different concentrations of magnesium ion [Mg2+], with color correspondence to 1 mM in green, 2 mM in orange, 3 mM in rosy color, and 4 mM in blue; with 20°C as constant temperature. **(f)** Tertiary structures of the eight aptamers were modeled at different concentrations of magnesium ion [Mg2+], with the same color correspondence. Molecular graphics, performed by PyMOL Molecular Graphics System, Version 2.0 Schrödinger, LLC [25].

In temperature as a focal point, Figure 4(a), we can agree to a constant behavior for most all the aptamer sequences. Still, it doesn’t mean that we can consider a stable parameter, so that reflects the necessity of *in silico* analysis, Figure 4(b) displays the tertiary structure when clearly TESS3 comes out as the most stable against temperature changes.

For ionic conditions, we set the temperature at 20° C, since we consider as an experimental parameter and to focus on the evaluation of the sodium ion (Na^+^) which also gives us an approach to other ions which is considered equivalent, like Li^+^, K^+^, and NH ^+^, Figure 4(c) and 4(d). There is a markable behavior for most all the aptamer sequences and great stability for TESS3. The evaluation of the magnesium ion (Mg^2+^) which also gives us an approach to another ion that is considered equivalent to Ca^2+^, Figure 4(e) and 4(f), has a markable behavior for all aptamer sequences and great stability for TESS3 [15].

Another point to evaluate sodium and magnesium ions is that the relation between the effects and the amount/concentration of these ions, cannot be considered a basic or direct effect, because only a specific range of concentration can hold the conformation while a low concentration conducts an unstable aptamer structure, and the counterpart, high concentration, cause electrostatic repulsions that interference on the aptamer binding [30,31]. *In vitro* methodologies allow us to determine a working concentration of both ions and also evaluate for each aptamer the ability to maintain its structure through the changed conditions. Preliminary experiments are performed on aptamers T5.1, T6, TESS1, TESS2, and TESS3 by considering the structural changes, showed in Figure 4, into a size (nanometers) change, values obtained with the technique of Dynamic Light Scattering performed by triplicate on the same temperatures and concentrations for *in silico* analysis. The average size of each evaluation is displayed in Figure 5.

**Figure 5.**
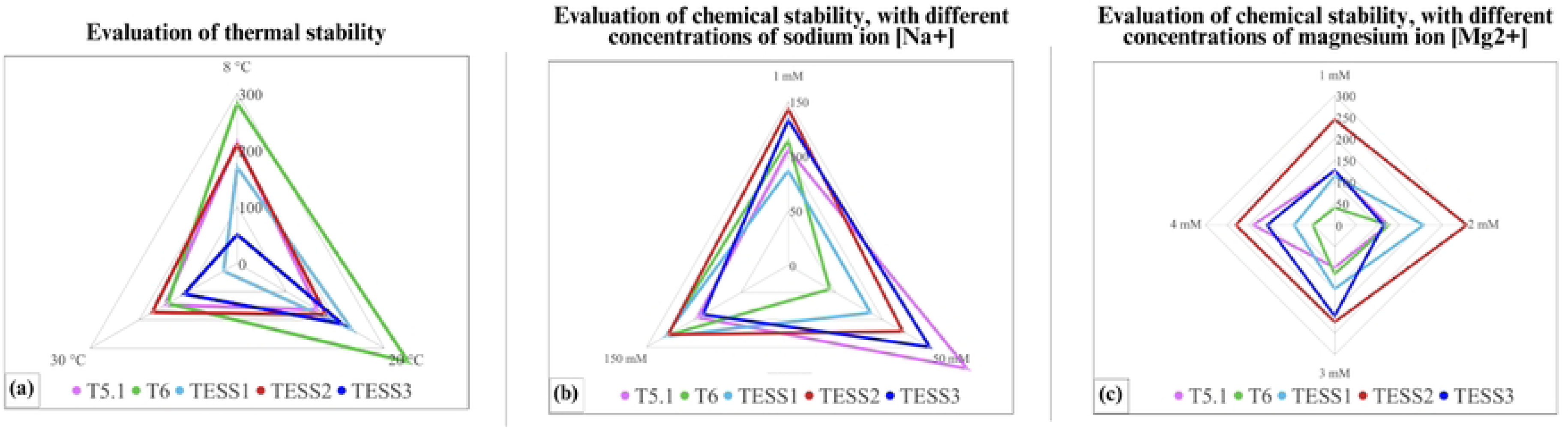
Area graph, according to average size (nm). In vitro analysis on Malvern Panalytical Zetasizer for the evaluation of physicochemical stability of aptamers. Color correspondence: T5.1 in violet, T6 in green, TESS1 in blue light, TESS2 in red, and TESS3 in dark blue. **(a)** Evaluation of thermal stability with the designation of axes at temperatures 8 °C, 20 °C, and 30 °C. **(b)** Evaluation of chemical stability by sodium ion [Na^+^] with the designation of axes at different concentrations of 1 mM, 50 mM, and 150 mM. **(c)** Evaluation of chemical stability by magnesium ion [Mg^2+^] with the designation of axes at different concentrations of 1 mM, 2 mM, 3 mM, and 4 mM.

The evaluation of the aptamer affinity is defined as molecular docking. Still, this analysis also provides information about the number and kind of interactions according to the previously tertiary structure designed. For molecular docking, we assume a flexible aptamer, and using AutoDock Vina as the program for molecular docking, the algorithm predicts the molecular interactions by sequence and structure databases [10,27].

Considering the tertiary structures of each aptamer at experimental conditions: magnesium ion concentration at 2 mM, sodium ion concentration at 50 mM, and a temperature of 20 ° C. The ΔG value, obtained on molecular docking, in the same way as in the construction of the secondary structure, we expect a negative value with a high magnitude to determine the most affine to the analyte (testosterone, androstenedione, and, testosterone undecanoate).

In a primary observation, TESS2, and TESS1 results were the most affine for the three analytes at any condition, TESS3 presented high affinity to testosterone and androstenedione at any condition, and T6 presented a great affinity for the three analytes in the presence of magnesium ion (2 mM).

Their structure and predicted interaction for aptamers apT5, T4, T5.1, T6, TESS2, and TESS3 are presented in Supplementary Material Figures B.1 to B.6. In most cases, the interactions are defined for conventional hydrogen bonds, carbon-hydrogen bonds, pi-alkyl, and pi-sigma, this kind of internal interactions are expected for each aptamer-analyte conjugations, also visualized in Supplementary Material Tables C.1 to C.6. Docking for TESS1 and P4G13 are presented since P4G13 is the aptamer with fewer interactions and demonstrates being unstable for the physicochemical aspect, and on the other hand TESS1 has great physicochemical stability and the highest affinity for the three evaluated analytes. Attending the information that provides the molecular docking of each aptamer, and getting a closer view and comparison, we present the following table, on blue rows are displayed the total interactions between the aptamer and the three analytes, followed by the total interaction for each type of bonding.

The column of affinity exhibits three categories according to the total of interactions in which the green arrow is the aptamer with higher affinity, the red arrow refers to aptamers with poor and unsatisfactory affinity, and the ones who have an intermediate number interaction are presented with a yellow script. A focused view of each analyte and their respective interactions are displayed in Supplementary Material, Table A.4. for testosterone, Table A.5. for androstenedione, and Table A.6. for testosterone undecanoate.

According to Table 2, we select only a few sequences to execute the *in vitro* analysis for an overview of the binding event, between testosterone and testosterone undecanoate as analytes, with the aim to compare the three affinity categories also expressed in Table 2. Employing well-established concentrations of analyte solutions that interact with the aptamers T5,1, T6, TESS1, TESS2, and TESS3, which are already coupled to gold nanoparticles (AuNPs)for its immobilization and separate, allows us to assess for a relative capture quantity. The results are presented as percentages in Figure 7, providing a visual representation of the capture efficiency for each aptamer.

**Figure 6.**
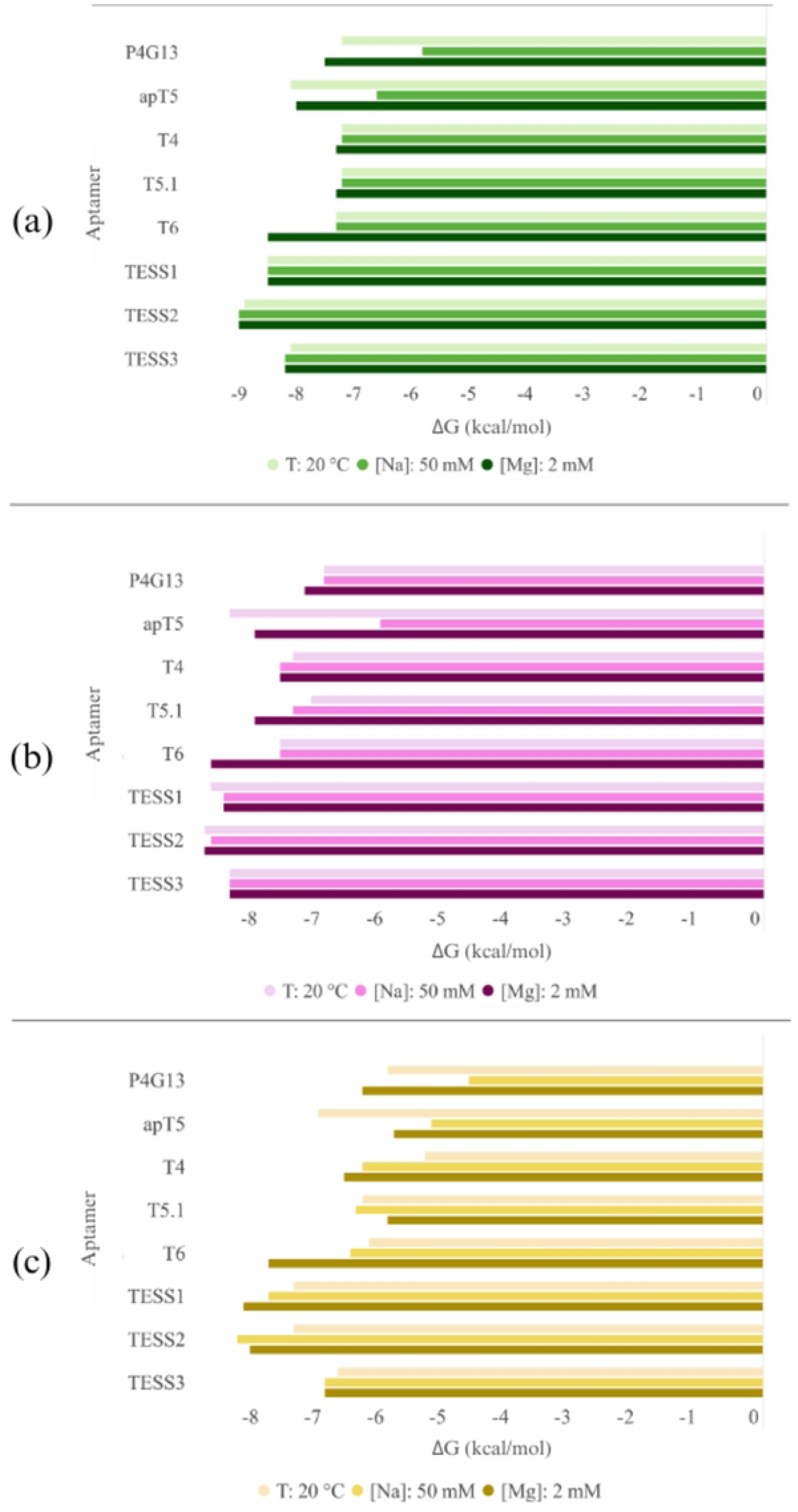
Evaluation of affinity in experimental conditions. Temperature of 20° C, sodium ion concentration of 50 Mm, and magnesium ion concentration of 2 mM. ΔG values obtained in the performance of molecular docking, Table A.3. (a) Evaluation of affinity with testosterone as the target. (b) Evaluation of affinity with androstenedione as the target. (c) Evaluation of affinity with testosterone undecanoate as the target.

**Figure 7.**
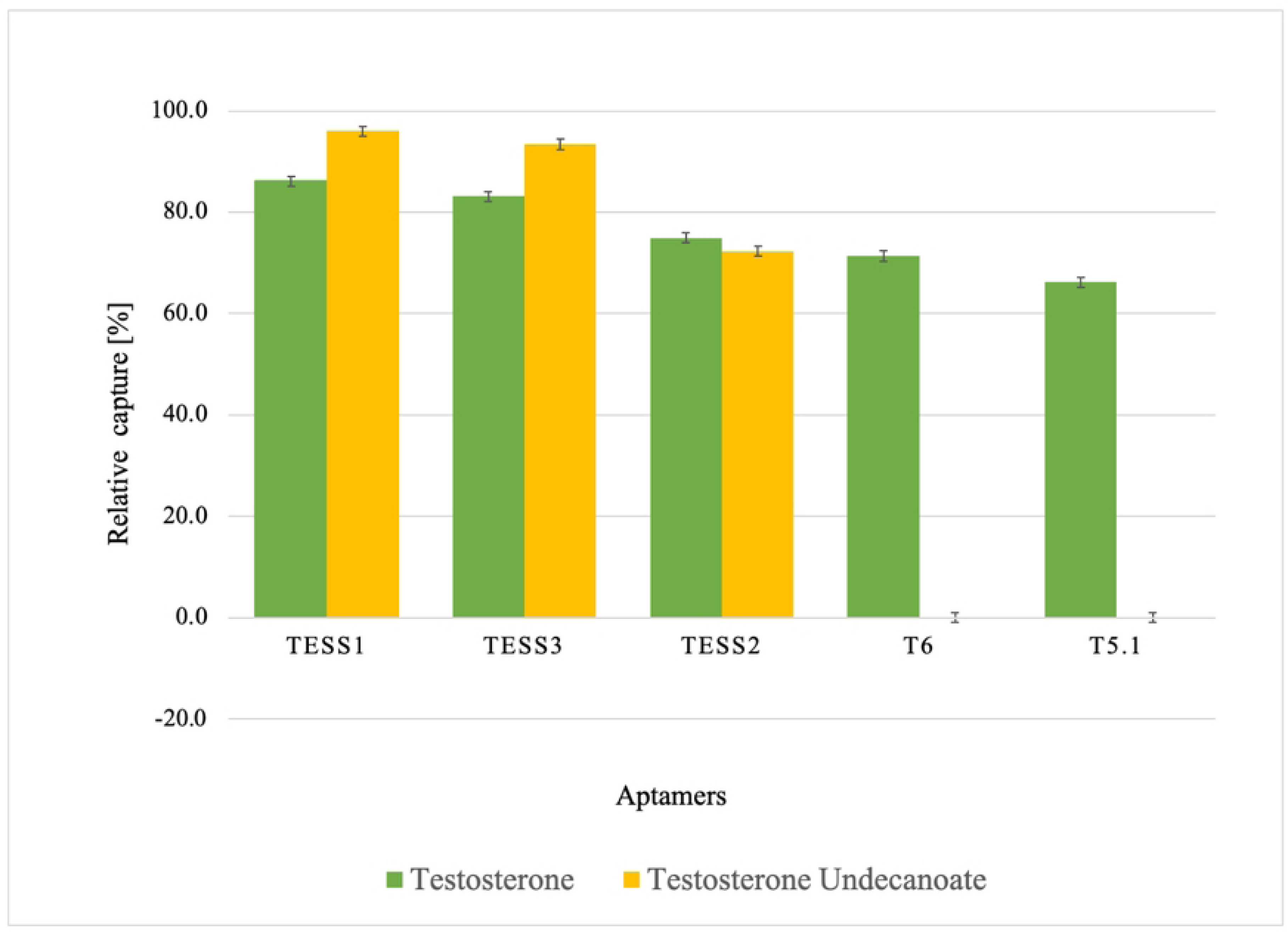
Evaluation of affinity. *In vitro* relative capture experiments for aptamers T5,1, T6, TESS1, TESS2, and TESS3, in experimental conditions of 20 °C and 50 mM of ion sodium concentrations. The average with a standard deviation of 3 replicates is presented.

**Table 2.**
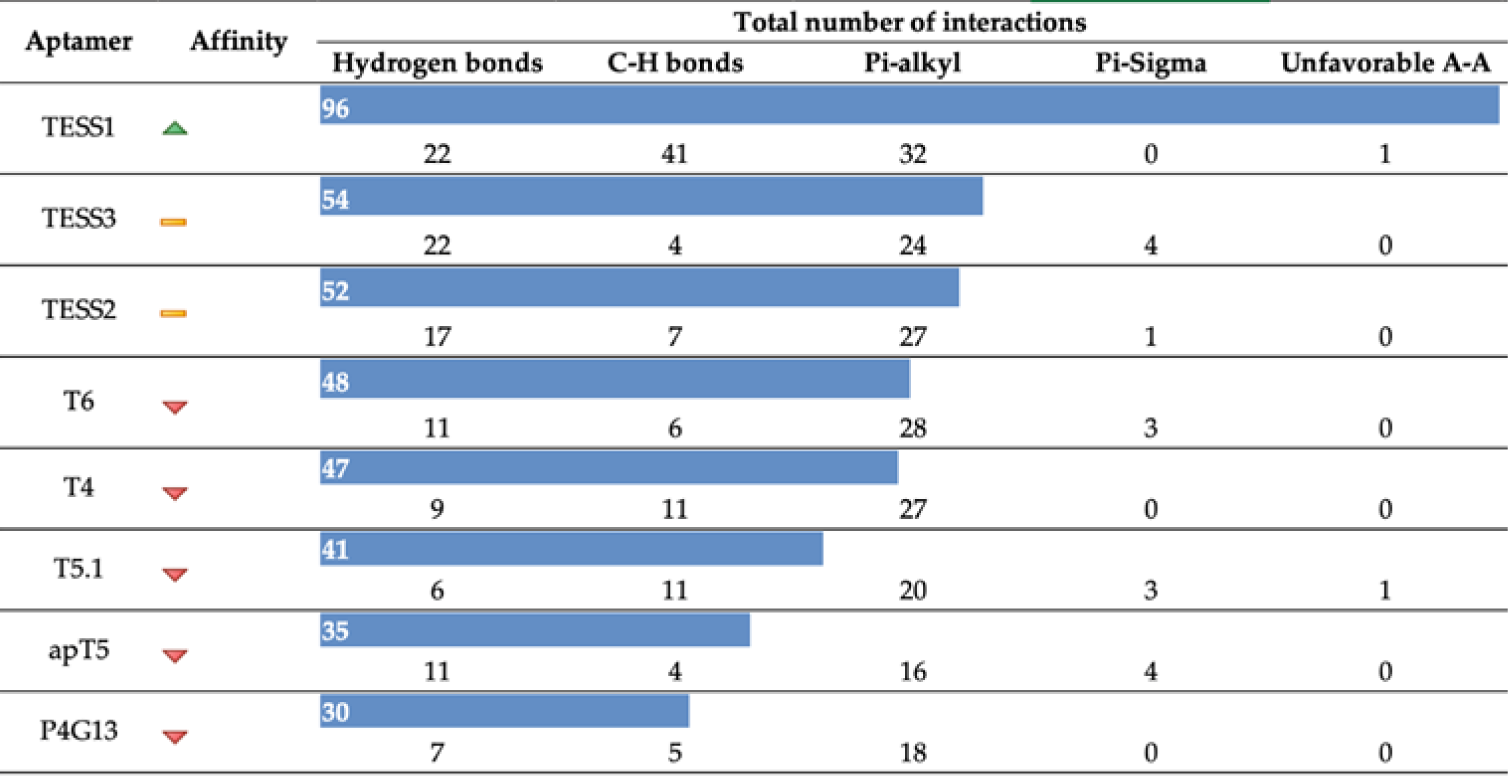
Summary of molecular docking and their interactions.

The correlation between Table 2 and Figure 7 underscores the comprehensive insight provided by *in silico* analysis into the experimental behavior. Furthermore, supports the assertion that the ability of aptamers to modify their three-dimensional conformation is influenced by environmental conditions. This dynamic conformational shift impacts their affinity with the analyte.

Aptamer TESS1 has the greatest affinity for the three analytes with a total of 96 interactions, displayed in Figure 8, where carbon-hydrogen bonds are the most prevalent followed by pi-alkyl. In Table 3, we can see that present more concurrent residues even with the change of the conditions, for testosterone T (4), C (5), C (10), and C (46); for androstenedione A (9) and C (10); for testosterone undecanoate G (7), G (11), T (42), and C (46). The relative capture experiment, Figure 7, points TESS1 as the aptamer with higher affinity for testosterone (86.13 %) and their synthetic analog testosterone undecanoate (96.01 %), results consistent with Table 2, indicated as the aptamer with higher affinity.

**Figure 8.**
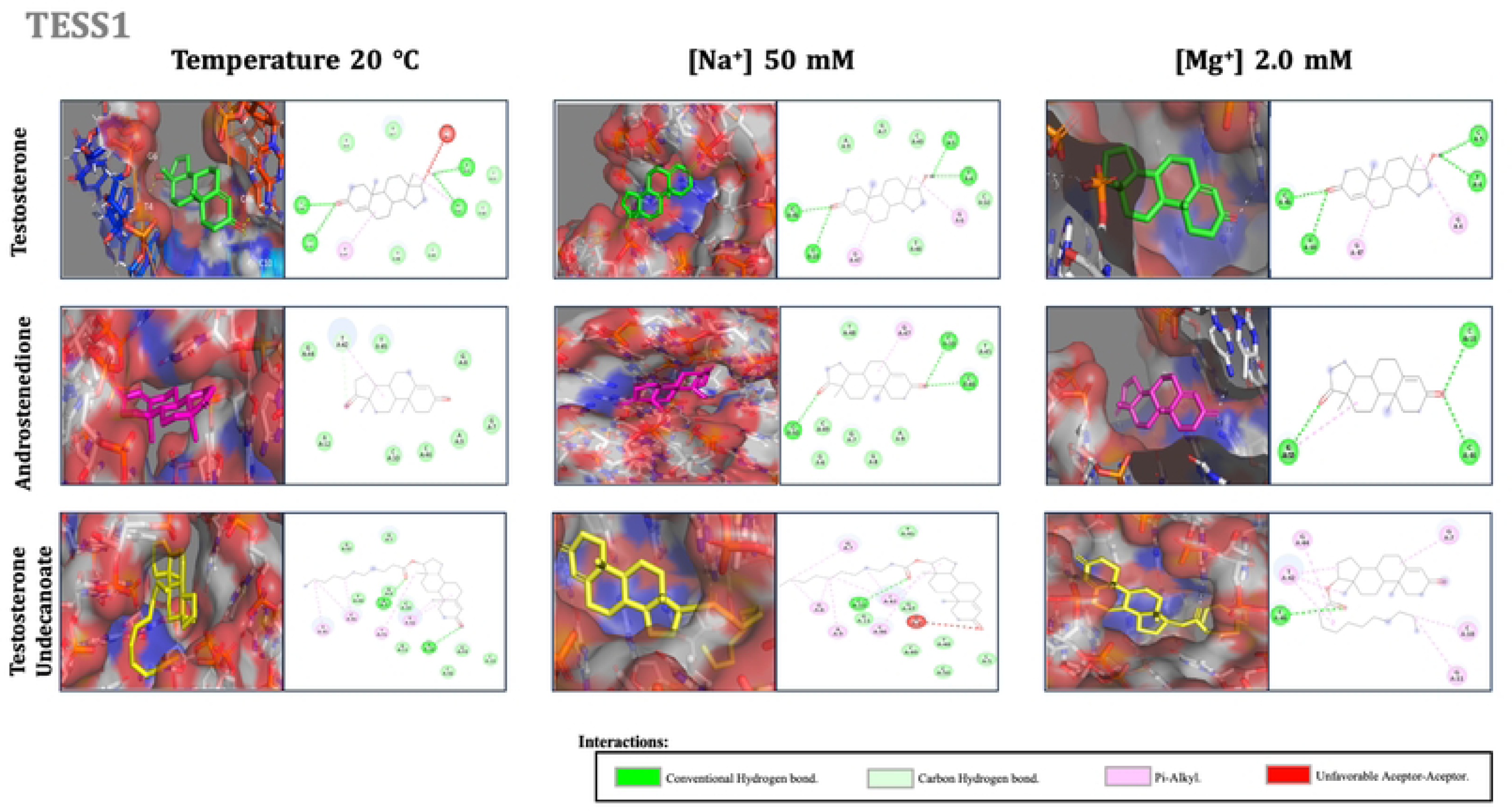
Molecular docking for TESS1. The color corresponds to the type of interactions, conventional hydrogen bonds in fluorescent green, carbon-hydrogen bonds in light green, pi-alkyl bonds in pink, and unfavorable aceptor-aceptor bonds in red. To visualize all aptamer–target interactions we develop the images on PyMOL Molecular Graphics System, Version 2.0 Schrödinger, LLC, for 3D structure visualization and BIOVIA, Dassault Systèmes, Discovery Studio Visualizer, v 21.1.0.20298, San Diego, for the images of 2D structures [25,28].

**Table 3.**
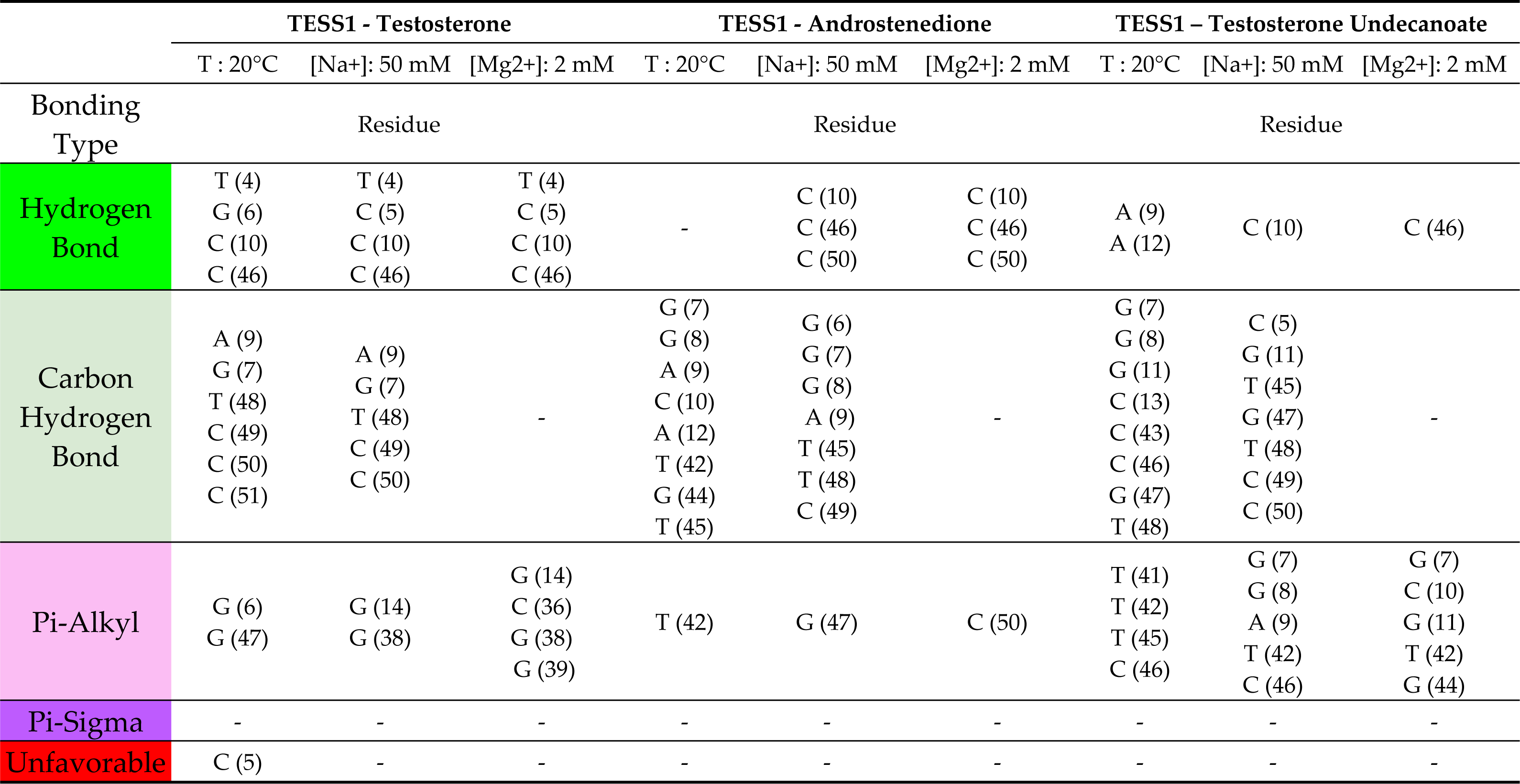
Interactions between aptamer TESS1 and analytes/ligands (testosterone, androstenedione, and testosterone undecanoate) are displayed in Figure 8.

TESS3 presents a total of 54 interactions distributed in four different interactions, where pi-alkyl and hydrogen bonds are the most common bonds since TESS3 demonstrated the highest stability at any condition. It is reasonable to visualize in Figure B.1, and Table C.1 more concurrent residues for all analytes. For testosterone G (7), G (8), and A (43); for androstenedione G (7), and G (10); for testosterone undecanoate G (7), G (10), A (12), T (42), and A (43). The number of interactions is higher for testosterone and testosterone undecanoate which means a higher affinity to both analytes. The last was corroborated by a relative capture experiment to get a percentage of the total amount of testosterone undecanoate binding or “capture” by the TESS3 in the same conditions of temperature and sodium ion concentration as their molecular docking, getting an 83.11 % binding with testosterone and a 93.42 % binding with testosterone undecanoate.

According to Figure B.2., TESS2 has a total of 52 interactions with pi-alkyl as the most common interaction with high affinity to testosterone and testosterone undecanoate. In Table C.2, we can visualize that the three analytes have G (7) as the frequent residue, testosterone has G (47), and androstenedione has C (46) and C (50). TESS2 in general differs only for two total interactions with TESS3, in Table 2, but in Figure 7, shows less affinity. The individual number of interactions for each analyte inverts the order between TESS2 and T6, Table A.4, for testosterone so the results are consistent with the *in silico* analysis, reducing their affinity also the absence of frequent residue impact on their relative capture (74.97 %). For testosterone undecanoate the position of affinity according to number interaction remains constant Table 2 and A.6, with a relative capture of 72.39 %.

According to Figure B.3, and Table C.3, T6 has a total of 48 interactions where more than a half are pi-alkyl and the highest affinity with testosterone with three concurrent residues C (36), G (38), and G (39); their residues have a similar presence with androstenedione that also has C (36) as a concurrent residue through the variation of conditions. In Table 2, T6 is the first aptamer with unsatisfactory affinity, they present less frequent residues as the aptamers who have intermediate affinity on a relative capture experiment of T6 at the same conditions of temperature and sodium ion concentration as their molecular docking, we obtain a 0.07 % binding with testosterone undecanoate. But as we already mentioned, testosterone presents an intermediate affinity, Table A.4, the reason for the percentage of relative capture of 71.33 % very close to TESS2.

According to Figure B.4 and their constructed Table C.4, we can find a total of 47 interactions for aptamer T4 distributed in three types of interactions being more than a half pi-alkyl bond. Also, T4 has more interactions meaning a highest affinity for testosterone undecanoate. There is no concurrent residue between analytes, but we find residues whose presence is not affected by the change of conditions, for testosterone G (13); for androstenedione G (4), A (5), and G (52); for testosterone undecanoate A (11), G (12), G (13), and C (24).

Aptamer T5.1 displays a total of 41 interactions, as we can visualize in Figure B.5, the most prevalent bonds are pi-alkyl and carbon-hydrogen bonds. In Table C.5, we can visualize a few concurrent residues, for testosterone G (38), for androstenedione G (13), and G (14). We can also see that there are no concurrent residues for testosterone undecanoate, so applying conditions has a negative effect on the development of interactions. It is also demonstrated with an unsatisfactory affinity with 66.20 % testosterone and 0.05 % testosterone undecanoate binding from their relative capture experiment, the lowest values among all the conducted experiments.

Aptamer apT5 has a total of 35 interactions, which are shown in Figure B.6, Table C.6 Pi-alkyl and hydrogen bonds are the prominent interactions, for all conditions and for different analytes/ligands. The most coincident residue is C (42) and it is presented in almost all kinds of interaction. Another important aspect in relation to the number of interactions, which is higher in testosterone followed by testosterone undecanoate and androstenedione, is that apT5 aptamer has more affinity to testosterone.

P4G13 has a total of 30 interactions but only exhibits three kinds of interactions: hydrogen bonds, carbon-hydrogen bonds, and pi-alkyl, Table 4 Pi-alkyl is the prominent interaction, and C (78) is the residue with the highest presence for all conditions and for different analytes/ligands. Furthermore, the interactions between testosterone and androstenedione are more related to each other than with testosterone undecanoate, which also turns out to be the analyte with the highest number of interactions, Figure 9.

**Figure 9.**
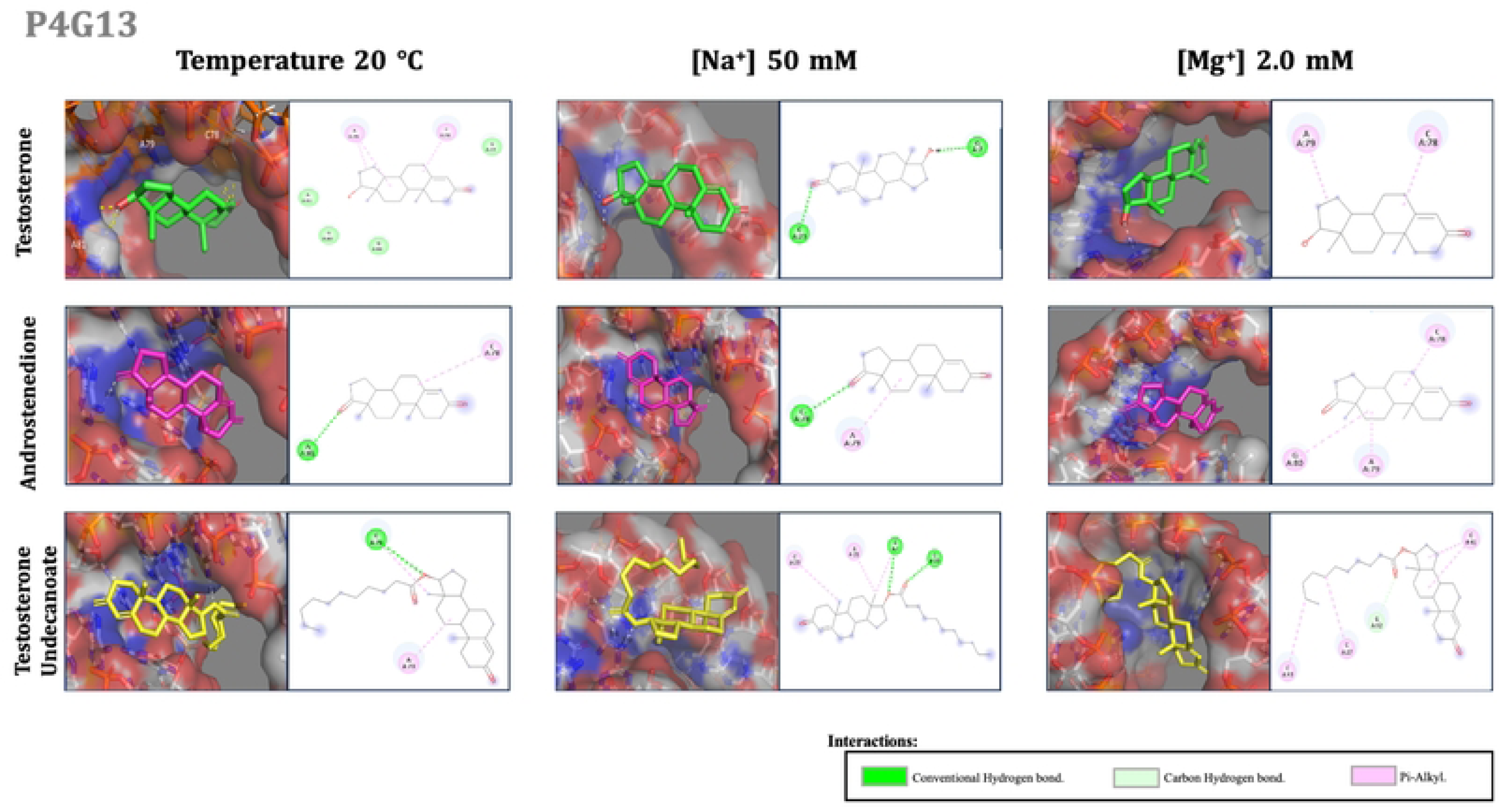
Molecular docking for P4G13. The color corresponds to the type of interactions, conventional hydrogen bonds in fluorescent green, carbon-hydrogen bonds in light green, and pi-alkyl bonds in pink. To visualize all aptamer–target interactions we develop the images on PyMOL Molecular Graphics System, Version 2.0 Schrödinger, LLC, for 3D structure visualization and BIOVIA, Dassault Systèmes, Discovery Studio Visualizer, v 21.1.0.20298, San Diego, for the images of 2D structures [25,28].

**Table 4.** Interactions between aptamer P4G13 and analytes/ligands (testosterone, androstenedione, and testosterone undecanoate displayed in Figure 9.

|  | P4G13 - Testosterone |  |  | P4G13 - Androstenedione |  |  | P4G13 – Testosterone Undecanoate |  |  |
| --- | --- | --- | --- | --- | --- | --- | --- | --- | --- |
|  | T : 20°C | [Na<sup>+</sup>]: 50 mM | [Mg<sup>2+</sup>]: 2 mM | T : 20°C | [Na<sup>+</sup>]: 50 mM | [Mg<sup>2+</sup>]: 2 mM | T : 20°C | [Na<sup>+</sup>]: 50 mM | [Mg<sup>2+</sup>]: 2 mM |
| Bonding Type | Residue |  |  | Residue |  |  | Residue |  |  |
| Hydrogen Bond | - | G (7)<br>C (23) | - | A (81) | C (78) | - | C (78) | G (7)<br>C (20) | - |
| Carbon Hydrogen Bond | G (77)<br>G (80)<br>A (81)<br>G (88) | - | - | - | - | - | - | - | G (42) |
| Pi-Alkyl | C (78)<br>A (79) | - | C (78)<br>A (79) | C (78) | A (79) | C (78)<br>A (79)<br>G (80) | C (78)<br>A (79) | G (7)<br>A (21)<br>C (23) | C (27)<br>C (41)<br>C (43) |
| Pi-Sigma | - | - | - | - | - | - | - | - | - |
| Unfavorable A - A | - | - | - | - | - | - | - | - | - |

## 4. Discussion

According to the results, we obtained useful information to select the optimal and rational aptamer or aptamer sequences for the design of a biosensor for testosterone detection.

The trending behavior of each aptamer was constant, and a comparison of the ΔG values was made in Figure 3, (Table A.1 & Table A.2), we can establish that magnesium ions have a greater effect than sodium ions since the minimal change in concentration increases their stability with little effect over the structural conformation. Also, the same values presented in Figure 1, (Table A.1 & Table A.2), are concentrated on a “database” in Minitab to perform a multiple regression analysis to show the relative contribution of independent variables (temperature, sodium, and magnesium ions) while controlling the others by the significance or non-significance relations between the response/dependent variable (ΔG values) [29]. For the multiple regression analysis, we ensured model adequacy by evaluating the equality of variances for each independent variable, using the Bartletts Method. Additionally, we assessed the normal distribution of residuals using the Anderson-Darling Normative Test. Finally, we implemented cross-validation as an integral component of the analysis, which shows a general R-squad of 99 %, for all the aptamers.

Statistical findings substantiate the pronounced impact on the stability (characteristic evaluated by ΔG values) from temperature, as the variable with a higher significative effect, followed by magnesium ions, which also exhibit a superior interaction intensity compared to sodium ions. Since the latter, manifests a low significant effect and demonstrates multicollinearity, indicative of an interference from other variables.

The thermal stability was evaluated primarily at a general room temperature (20°C) and from their lower and higher values. The aptamer’s behavior showed that as the temperature increases the structure is more unstable, getting closer to ΔG values equal to zero, that is explained because at higher temperatures the oligonucleotides tend to break the hydrogen bonds and possible electrostatic interactions between nucleotide bases. In Figure 4(a), apT5, and P4G13 show the highest (positive or near to zero) ΔG values, and also significant changes in their structures, which decreased the possibility of adapting to the conditions in which the tests were executed. The aptamers whose lowest (negative) values are the most stable are also demonstrated in Figure 4(b), like TESS3 and TESS2 which have no structural changes, aptamers and TESS1 which has minimal conformational changes, between temperatures of 8°C and 20°C, these three aptamers have in common a stem-loop as their principal structural conformation. T4, T5.1, and T6 also have minimal conformational changes, between temperatures of 8°C and 20°C, where T6 and T5.1 present various structures, like a hairpin, helix, and internal loop, but the majority part of the sequences corresponds to a multi-branched loop, similarly, T4 present different structures like a hairpin, helix, bulge, internal loop, and multi-branched loop. *In vitro* methodology, is consistent with the statistical analyses concerning the fact that temperature exerts a greater effect on the structure and therefore stability of the aptamer, which is reflected in the change in size (nm) for the five aptamers T5.1, T6, TESS1, TESS2 and TESS3 evaluated experimentally, Figure 5. Likewise, it is important to mention that T5.1, quantitatively appears to have a certain stability but by carrying out the experiment it was observed that the sample had many fluctuations of significantly larger sizes, therefore, it is inferred that the temperature change is what generates a denaturation effect on the aptamer.

When looking at chemical stability, we evaluate ions commonly present in buffers, sodium ion starts from a worked concentration of 50 mM and modify it into a higher and lower value, and magnesium ion performs in general concentrations, according to the literature, since it is not an obligate required ion for our future worked solution on the binding event [30,31]. For sodium ion concentration, [Na^+^], the ΔG value (kcal/mol) decreases as the sodium concentration increases in response to the capacity of sodium as a stabilizing cation for nucleic acids, reducing the solubility that surrounds the sequence, allowing it to generate a greater number of intra-interactions. From Figure 4(c) and 4(d), we can observe both in the surface graph and in the modeling of tertiary structure that the aptamers apT5 and, P4G13, are not stable and present notable structural changes. T4 and T6 present great structural changes despite having a very low value, being modeled at 150 mM, this results from the conformation of the nucleotide bases of the T4 sequence in addition to being a consequence of the concentration of sodium, since this sequence is presented more compactly, forming a greater number of internal loops between itself. T5.1 has similar conformation structures with little difference at the end of the sequences adding a hairpin structure. On the contrary, the aptamer TESS3 presents values that have the quality of preserving its molecular structure, as a stem-loop, against any change. TESS1 and TESS2 present slight structural changes such as forks, internal loops, and hairpins, and a stable structure between concentrations of 50mM to 150mM, being suitable for future experimental phases.

The disparities between *in silico* and *in vitro* experiments across various aptamers arise from distinctions in their primary structure, sequence, and length to their 3D structural conformation. *In vitro* analyses, as *in silico* analysis, offer insights into the behavior and stability of aptamers, but only *in vitro* analysis sheds light on the concentrations in which the studied aptamer maintains structural integrity. This is crucial because ion concentrations can impact the aptamer’s structure, and consequently diminish or interfere with its binding capability. In examining and comparing the sizes of aptamers T5.1, T6, TESS1, TESS2 and TESSS3 with their respective controls (dilution under any sodium and magnesium concentrations at room temperature), the *in silico* expectation of TESS3 greater stability over TESS1 is inverse, but don’t have negative results on the use of both aptamers since most of their size variations fall within the standard deviation of control measurements, assuming an absence of structural changes. Notably, TESS1 emerges as the most stable and functional aptamer, demonstrating viability across sodium concentrations ranging from 1 to 150 mM. On the other hand, TESS3 is deemed safe for operation within a sodium concentration range of 1 to 50 mM. *In silico*, and *in vitro* result agrees for aptamer TESS2 with greater stability at higher temperatures (30 °C), at any sodium concentration TESS2 remains stable since their size variations fall within the standard deviation of control measurements, Figure 5 (b), but for magnesium concentrations aptamer TESS2 *in silico* evaluation, Figure 4 (c) even the change in different magnesium concentrations can assume minimal modification it demonstrates a complete structural change for the aptamer compared with their conformation at others conditions and more important in an optimal perspective, so as it has been demonstrated *in vitro* evaluation, Figure 5 (c), with a big graph area. For T6, it is challenging to establish a range of concentrations, and inferences from *in silico* simulations, Figures 5(c) and 5(d), of its tertiary structure at different sodium concentrations, only certain regions change while other regions share their structure within different concentrations but without being consistent. T5.1 has slight changes and a certain structural pattern was observed, in Figures 4(c) and 4(d), however experimentally structural data reveal significant size variations, rendering T5.1 unsuitable for our biosensor project. Specifically, because a minimum sodium concentration of 50 mM is required on the binding event with the analyte.

Evaluating the aptamers in concentrations of magnesium ion [Mg^2+^], the ΔG value (kcal/mol) decreases as the concentration of magnesium increases, this phenomenon occurs because the magnesium ion has the characteristic of joining with the deoxyribose and forming complexes, generation an evident effect for the aptamer conformation since its develop more structural change (internal loops, harpins and bungle) comparing with the structures obtained at sodium ion concentrations and temperature changes. For the magnesium ion only the aptamer apT5 presents a noticeable structural change as an extended helix in 4.0 mM, while TESS3 and P4G13 are stable at any concentration. The rest of the aptamers show slight differences that are related more to their compaction than structural changes through the increases in magnesium ion concentration; T4 and TESS1 present a different conformation when the concentration changes from 2 mM to 3 mM. TESS2, T5.1, and T6 display a different conformation only for 1 mM being stable at rest and usable for future experimentation phases. The *in silico* structural conformations of aptamers through magnesium ion give us a little perspective of the concentrations on the aptamer’s present stability but *in vitro* measurements establish, that the size variations of TESS3 fall within the standard deviation of control measurements, assuming an absence of structural changes, same for TESS1, being both aptamers functional on a range concentration of 1 to 4 mM. While T6 is only useful at concentrations of 2 to 3 mM similar to their *in silico* observation. And T5.1 is useful at concentrations of 1 to 2 mM.

For physio-chemical evaluation, we agree that TESS1 and TESS3 are stable and can be considered as the principal aptamers to use as a biosensing system. Only from an *in silico* point of view, we can consider great stability from TESS2. Also from an *in silico* approach, we consider neutral T5.1, T6, and T4 since they have some changes in their conformation but *in vitro* experiments show limitations of aptamer T5.1 and T6 on maintenance of their stability at a minimum work concentration of 50 mM, for T4 it’s needed to carry out experimental methodology.

Finally, the unstable ones, according to their *in silico* analysis, are apT5 and P4G13 which visually present significant changes being rejected from the *in silico* analysis according to Figures 3 and 4. Also, most “stable” aptamers present conformational arrangements of a multi-branched loop as the principal structure, and some hairpins and loops and the ones that are less, and the “un-stable” aptamers present more helix structure than loops.

Concerning the docking molecular results, we selected the average conditions of experimental works, related to the *in silico* and *in vitro* physicochemical analysis: a temperature of 20°C, sodium ion concentration of 50 mM, and magnesium ion concentration of 2 mM. Figure 6, represents the general panorama of the behavior of each aptamer to observe some trends and the expected quantitative stability. Considering that a negative value represents a stable structure whose numerical magnitude is proportional to its stability.

Figures 8 and 9, and from supplementary material Figures B.1. to B.6., show molecular dockings under the three conditions mentioned and for three analytes: testosterone and two synthetic analogs, androstenedione, and testosterone undecanoate. As indicated in each figure the molecular graphs were generated by two platforms. The most noticeable difference would be the format of 3D visualization for PyMOL and 2D for Discovery Studio [25,28]. However, the main difference is that PyMOL allows us to observe the conformation of the aptamer before the conditions and before the analyte, formulating a kind of cavity or pseudo cavity, denominated as a binding site, although it is possible to visualize the hydrogen bonds between aptamer and analyte. Discovery Studio will be the platform to see all kinds of interactions and the residue involved in those interactions [25,28].

According to Table 2, we found that the aptamer P4G13 has interactions of hydrogen bonds, carbon-hydrogen bonds, and pi-alkyl bonds. In the same way for apT5 with the difference of having a pi-sigma bond. P4G13 and apT5 are the aptamers with fewer interactions, most are pi-alkyl interactions and without a prominent concurrence in relation to the residuals which is consistent with its stability evaluation. Having values of ΔG close to zero and/or positive and modifications in its three-dimensional structure altering the establishment of interactions, and there is a low affinity with androstenedione, Figure 6, with the presence of no more than 7 interactions for this analyte in Figure 9 and B.6.

Aptamers such as T4, T5.1, and T6, result in many interactions and are well distributed in the different types of interactions to a greater extent are pi-alkyl bonds, being a type of link characteristic of the complex aptamer-analyte. These aptamers presented more affinity to testosterone, followed by testosterone undecanoate, and at the end with minimal interactions for androstenedione. This can be explained because testosterone undecanoate is a version of esterified testosterone, and the change of residues between testosterone and testosterone undecanoate is due to the arrangement of the aliphatic chain within the binding site (pseudo cavity) that forms the aptamer in its tertiary structure, same reason why there is no prevalence and any residue for testosterone undecanoate with T5.1 and T6. The last two aptamers show almost null capture for testosterone undecanoate. But more than 50 % of capture for testosterone, from these results we can infer that T5.1 and T6 are very specific to testosterone.

TESS2 and TESS3 have a greater number of interactions concentrated in conventional hydrogen bonds and pi–alkyl bonds with more common residues between analytes and conditions. Only TESS2 shows a great affinity for the three analytes while TESS3 has minimum affinity for androstenedione. On an experimental and closer look at testosterone and testosterone undecanoate as analytes, the affinity is higher on TESS3 than TESS2, the same result that shows the molecular docking represented in Table 2 and Figure 7. TESS1 represents the aptamer with the highest affinity for any of the three analytes since it presents almost the double number of interactions in contrast to the other aptamers. Their principal kind of interactions are carbon-hydrogen bonds, followed by pi–alkyl bonds, and hydrogen; not only have a preferable distribution of the type of interactions but also have many residuals remaining as interaction targets. Also demonstrated by the relative capture experiment obtaining testosterone and testosterone undecanoate with the highest percentage. As we already mentioned TESS1 is one of the two principal aptamers physicochemical stable, and also presents the greatest capture for testosterone and its synthetic analog (testosterone undecanoate) in both perspectives (i*n silico* and *in vitro*). These results demonstrate the capacity and competence of TESS1 to be used as a biosensing system and easily coupled to a transducer or other platform, without reducing its affinity and specificity, being this characteristic the most desirable attribute of the receptor/capture molecule on a biosensor.

It’s important to mention that as a banned substance, testosterone and its analogs are only commerce in Mexico in pharmaceutical presentation, also controlled, the reason why androstenedione couldn’t be tested on the *in vitro* analysis.

Considering that the equilibrium dissociation constant (*Kd*) is used to assess, and rank order the strengths of bimolecular contacts and report binding affinity, we collected the dissociation constant (*Kd*) of the nine aptamers analyzed. It is worth mentioning that for T4 there are no reports of its *Kd*. For T6, T5.1, apT5, and p4G13 this value has only been estimated theoretically. The corresponding values are 800 nM (T6), 0.5 nM (T5.1), 4.0 +/- 5.8 nM (apT5), and 1.65 nM (P4G13) [18 - 20]. Subject to being theoretical values, it is observed that T5.1, apT5, and P4G13 have a high affinity for the target molecule. However, experimental determination of their *Kd* has been made for 3 aptamers, 80 nM for TESS1, 1700 nM for TESS2, and 700 nM for TESS3 [21]. The results obtained in this study, by molecular docking, coincide with pointing out TESS1 as the aptamer with the highest affinity, in the case of those aptamers for which the *Kd* has been determined experimentally.

In summary, the *in silico* or simulation analyses provide useful information regarding the selection of the best aptamer candidate for their use on biosensors, by ascertaining the ability to preserve the nature and structure of each of the proposed aptamer sequences. However, it is imperative to evaluate the affinity with analytes, although the number and type of interactions, referring type, to which it is expected. These interactions do not correspond to simple covalent bonds, they are multiple bonds of different types. A pattern of repetition of residues capable of interacting between each aptamer-analyte complex is also observed. The number and richness of bonds allow us to confirm the affinity of each aptamer for the corresponding analyte. In this study and from an *in silico* perspective the aptamer with the best evaluation of stability and interaction with testosterone and the two analogs used was TESS1. From an *in vitro* perspective and with a capture relative experimentation, Figure 5, for two aptamers considered as stable and with intermediate stability, TESS21 and TESS3 achieve a good affinity, principal TESS1 (86.13 % for testosterone and 96.01 % for testosterone undecanoate) that also demonstrate to be stable on the set conditions of temperature, sodium ion, and magnesium ion. Opposite, aptamers T5.1 and T6 that not only cataloged with poor or unsatisfactory affinity by their molecular docking beside their *in silico* analysis present structural changes and *in vitro* analysis shows instability on temperature and primarily on the minimum concentration of 50 mM of ion sodium that assumes have remarkable negative effects on their binding with testosterone undecanoate experiments.

## 5. Conclusion

The application of bioinformatics tools in relation to aptamers is commonly related mostly as an auxiliary tool in SELEX. However, the usefulness of the selection of the sequences obtained is equally important since, as it was observed in the present writing some of the sequences are not viable for experimental developments and/or certain types of uses. While the analysis of interactions is useful since the best candidate is sought, it is true that all sequences are expected to be related and capture the analyte, but the nature of each sequence, as well as the tertiary structure that is built, will be key to establishing this affinity. Likewise, experimental results support the *in silico* analysis, indicating the utility and truthfulness of this computational approach.

We discarded aptamers that did not possess thermal stability, such as the aptamer P4G13, and/or stability in the ionic plug, as apT5 in the compatible conditions used to run the tests. Performing molecular docking has little affinity with respect to the rest of aptamers. The rest of the aptamers present stability under experimental conditions and affinity for analytes. Those that stand out from the rest would be TESS3 which presents great stability against any change either thermal or chemical, but nevertheless, the aptamer with greater stability and equal level of affinity is TESS1, being proposed then as the best candidate for future purposes stipulated in the introduction section. The results obtained and the conclusions presented according to them are limited to the *in silico* and *in vitro* evaluation reported in this study.

## Appendix

Displayed in Supplementary data that can be found online at https://doi.org/XXX/XXXX

## Author Contributions

Conceptualization, A.M.B., A.A.P., and A.L.T.H.; methodology, A.M.B., A.A.P., and A.L.T.H; software, A.M.B.; validation, A.M.B., A.A.P., and A.L.T.H.; formal analysis, A.M.B.; investigation, A.M.B., A.A.P., and A.L.T.H.; resources, A.M.B., C.A.G.A, R.F.R; data curation, A.M.B.; writing—original draft preparation, A.M.B.; writing—review and editing, A.M.B., A.A.P, A.L.T.H., C.A.G.A., and R.F.R.; visualization, A.A.P, A.L.T.H., C.A.G.A., and R.F.R.; supervision, A.M.B., A.A.P., and A.L.T.H; project administration, A.M.B., A.A.P., and A.L.T.H; funding acquisition, A.M.B. All authors have read and agreed to the published version of the manuscript.

## Institutional Review Board Statement

Not applicable.

## Informed Consent Statement

Not applicable.

## Data Availability Statement

Not applicable.

## Acknowledgments

A.M.B. gratefully acknowledges the scholarship from CONAHCyT to pursue her postgraduate studies (No. CVU: 1065626). Special recognition and thanks to Dr. Rigel Valentín Gómez Acata for his teaching and advice on the construction of the charts developed on MATLAB.

